# Identification of a common deletion in the spike protein of SARS-CoV-2

**DOI:** 10.1101/2020.03.31.015941

**Authors:** Zhe Liu, Huanying Zheng, Runyu Yuan, Mingyue Li, Huifang Lin, Jingju Peng, Qianlin Xiong, Jiufeng Sun, Baisheng Li, Jie Wu, Ruben J.G. Hulswit, Thomas A. Bowden, Andrew Rambaut, Nick Loman, Oliver G Pybus, Changwen Ke, Jing Lu

## Abstract

Two notable features have been identified in the SARS-CoV-2 genome: (1) the receptor binding domain of SARS-CoV-2; (2) a unique insertion of twelve nucleotide or four amino acids (PRRA) at the S1 and S2 boundary. For the first feature, the similar RBD identified in SARs-like virus from pangolin suggests the RBD in SARS-CoV-2 may already exist in animal host(s) before it transmitted into human. The left puzzle is the history and function of the insertion at S1/S2 boundary, which is uniquely identified in SARS-CoV-2. In this study, we identified two variants from the first Guangdong SARS-CoV-2 cell strain, with deletion mutations on polybasic cleavage site (PRRAR) and its flank sites. More extensive screening indicates the deletion at the flank sites of PRRAR could be detected in 3 of 68 clinical samples and half of 22 in vitro isolated viral strains. These data indicate (1) the deletion of QTQTN, at the flank of polybasic cleavage site, is likely benefit the SARS-CoV-2 replication or infection in vitro but under strong purification selection in vivo since it is rarely identified in clinical samples; (2) there could be a very efficient mechanism for deleting this region from viral genome as the variants losing 23585-23599 is commonly detected after two rounds of cell passage. The mechanistic explanation for this in vitro adaptation and in vivo purification processes (or reverse) that led to such genomic changes in SARS-CoV-2 requires further work. Nonetheless, this study has provided valuable clues to aid further investigation of spike protein function and virus evolution. The deletion mutation identified in vitro isolation should be also noted for current vaccine development.

## Introduction

SARS-CoV-2 is a novel coronavirus firstly identified at the end of December 2019^1^ but has caused a global pandemic of COVID-19^2^. Unlike the other two zoonotic coronaviruses SARS CoV-1 and MERS^3^, the genetic evolution history is mostly unknown for SARS-CoV-2. A recent analysis based on the genetic information and protein structure highlights there are two notable features in the SARS-CoV-2 genome:

1. the receptor binding domain (RBD) of SARS-CoV-2 is distinct from the most closely-related bat-origin SARs related virus (RaTG13) and is demonstrated to have a high affinity to human ACE2 receptor;
2. a unique insertion of 12 nucleotides (or four amino acids, PRRA) at the S1 and S2 boundary results in a polybasic (furin) cleavage site and three predicted O-linked glycans around the cleavage site^4^.

With respect to the first feature, the similar RBD identified in a SARs-like virus from a pangolin suggests that the RBD in SARS-CoV-2 may already exist in its potential animal host(s) before it transmitted into human^5^. The question remaining is the history and function of the insertion at the S1/S2 boundary, which is uniquely identified in SARS-CoV-2. The insertion of proline is predicted to result in three addition of O-linked glycans. The functional consequence of the polybasic cleavage site and O-linked glycans in SARS-CoV-2 is unknown. By sequencing the whole genome of SARS-CoV-2, we identified two variants having deletion mutations on polybasic cleavage site (PRRAR) and its flank sites. More extensive screening indicates the deletion at the flank sites of PRRAR have been frequently observed in cell isolated strains and could be verified by multiple sequencing methods.

### Identification of deletions in SARS-CoV-2 spike protein

The first COVID-19 clinical case in Guangdong was reported on 19^th^ January, with illness onset on 1^st^ January^6^. A BALF (Bronchoalveolar lavage fluid) sample from this patient was collected and inoculated into Vero-E6 cells. The cell-isolated viral strain was obtained after three rounds of passage. Multiple sequencing methods were used for whole genome sequencing and the validation of variants (Figure A), including multiplex-PCR with Miseq platform (PE150), direct CDNA sequencing in Nanopore platform and Sanger sequencing. After mapping to the SARS-CoV-2 reference genome (MN908947.3), we found there were two variants in cell-isolated viral strain with deletions at (1) 23585–23599, flanking the polybasic cleavage site, resulting in a QTQTN deletion in spike protein (one amino acid before the polybasic cleavage site) and (2) 23596–23617, including the polybasic cleavage site and the 6 nucleotides 5’ of the cleavage site, resulted in a NSPRRAR deletion that included the polybasic cleavage site (Figure A). To exclude the possible errors caused by PCR amplification, both of these two deletion variants were verified through direct cDNA sequencing on the ONT nanopore platform. Sanger sequencing with specific primers also identified heterozygous peaks with distinct double peaks starting at the position 23585 and triple peaks after that, highlighting the existence of multiple variants caused by the above two deletions (Figure B).

**Figure.**
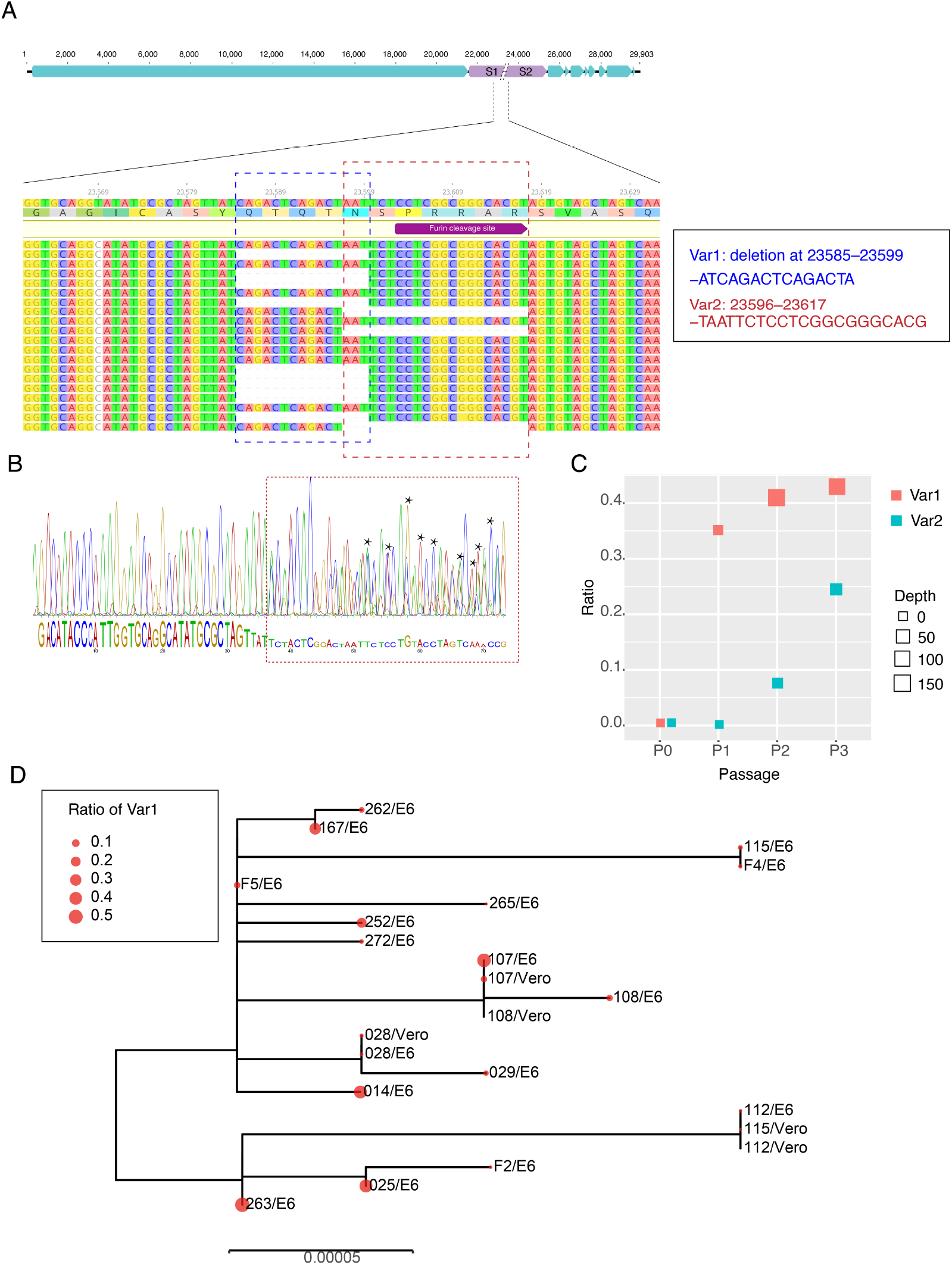
Deletion variants identified in SARS-CoV-2 cell strains. (A) High-throughput sequencing of the cell isolated strain (014) from the first SARS-CoV-2 patient (EPI 403934) in Guangdong, China. Representative reads mapping to the SARS-CoV-2 genome (MN908947.3 used as reference genome) showed two deletion variants. (B) Sanger sequencing of the 014 cell strains. The heterozygous peaks highlighted with a red box and the sites with distinct three peaks were marked with * (C) High-throughput sequencing showed the ratio of deletion variants in original clinical sample SF014 (P0) and cell strains after 3 rounds of cell passage (P1-3). (D) Phylogenetic tree of genome sequences of all 22 SARS-CoV-2 cell strains. The size of red dots is proportional to the ratio of Var1 (deletion at 23585–23599).

### The deletion is commonly identified in cell isolated strains

To investigate whether these deletions described above are random mutations occasionally identified in a strain or would commonly occur after cell passages, we performed whole genome sequencing on the other 21 SARS-CoV-2 viral strains collected after 2 rounds of cell passage in Vero-E6 or Vero cells (Supplemental Table). The corresponding original samples for these strains were collected between 19^th^ January and 28^th^ February 2020. Multiplex-PCR combined with the nanopore sequencing was used, following the general protocol as described in (https://artic.network/ncov-2019). The ARTIC pipeline was applied to trimmed primers and generated the bam files, which included all reads mapping to the SARS-CoV-2 reference genome (MN908947.3). Variant sites were called by using iVar^7^ with depth >=20 as a threshold. With this method, 10 of 21 cell isolate strains have different ratios of variants (>10%) with deletion at the flank of the polybasic cleavage site (deletion at 23585–23599) (Figure C). One has the variant with deletion on the polybasic cleavage site (deletion at 23596–23617). To find out whether the deletion on 23585–23599 was restricted in a specific genetic lineage, we next investigated the phylogenetic relationship of these strains and first 014 strain described above. As shown in Figure D, the strains with a relative higher ratio of this deletion were dispersed in the phylogenetic tree suggesting the deletion mutation was not restricted to a specific genetic lineage of SARS-CoV-2 viruses.

### Screening for deletion variants in original clinical samples

To identify whether these deletions also occurred in original clinical samples, we screened the high through-put sequencing data from 149 clinical samples, which collected between 6^th^ February and 20^th^ March in Guangdong, China. These samples were sequenced as by using multiplex PCR combined with nanopore sequencing. There were 68 SARS-CoV-2 genomes with sequencing average depth >=20 at the sites neighboring 23585. As shown in Table 1, the variants with the deletion at 23585-23599 were found in 3 (6%) of clinical samples with ratios ranging from 8.8–32.8% indicating this deletion may also occur in vivo infections even though the rate was extremely low compared to the results from in vitro (Figure D). To date, there are no genome sequences deposited in public dataset having this deletion. However, this did not mean this variant did not exist in currently released sequences since most of the variants with a lower ratio would be discarded when generating the final consensus sequences.

**Table 1:** Deletion variant (23585–23599) identified in clinical samples

| Samples | REF_depth | ALT_depth | Del Variant Ratio |
| --- | --- | --- | --- |
| 20SF5645 | 104 | 25 | 19.4% |
| ST-N3-D | 82 | 8 | 8.8% |
| SZ-N16-D | 256 | 125 | 32.8% |

The spike protein of coronaviruses plays a important role in viral infectivity, transmissibility and, antigenicity. Therefore, the genetic character of the spike protein in SARs-CoV-2 would shed light on its origin and evolution. For SARs-CoV-1, strong positive selection has been identified in the spike coding sequence^8^ and deletions in the other gene segment^9^ at the early stage but not the late stage of the epidemic, suggesting the adaptive pressures operated on the SARS-CoV-1 genome at the beginning of the epidemic. This result also indicates the SARs-CoV-1 may not well established in the human population at the early stage when it first transmitted from an intermediate animal host. For SARs-CoV-2, the virus presents high infectivity and efficient transmission capability among the human population since it is firstly identified^1^. Genetic changes related with viral fitness of SARs-CoV-2 require further epidemiological investigation and functional experiments.

Here, we use different sequencing methods to identify and verify a deletion at sites flanking the polybasic cleavage site. The deletion variants could be detected from 3 in 68 clinical samples, but half of 22 in vitro isolated viral strains tested in this study. These data indicate (1) the deletion of QTQTN may benefit SARS-CoV-2 replication or infection *in vitro* (Vero-E6 cell) but is likely to be under strong purification selection *in vivo* since it is rarely identified in clinical samples and (2) there could be an efficient mechanism for deleting this region from the viral genome, as the variants with the 23585–23599 deletion are commonly detected after two rounds of cell passage. Notably, a recently reported SARs-like strain RmYN02, which is phylogenetically related to a SARS-CoV-2, also has a deletion at the QTQT site^10^. This raises another possible scenario, which is that SARS-CoV-2-like viruses in animals may not have QTQTN in their spike protein and a variant with this insertion occurred upon virus transmission into humans. The mechanistic explanation and functional significance of these genomic changes in SARS-CoV-2 requires further work. Nonetheless, this study has provided valuable clues to aid further investigation of this remarkable evolutionary tale. The deletion mutation identified *in vitro* should be also noted for current vaccine development.

## Data Availability

Metagenomic sequencing, multiplex PCR sequencing and cDNA direct sequencing data after mapping to SARs-COV-2 reference genome (MN908947.3) have been deposited in the Genome Sequence Archive^11^ in BIG Data Center^12^, Beijing Institute of Genomics (BIG), Chinese Academy of Sciences, under project accession numbers CRA002500, publicly accessible at https://bigd.big.ac.cn/gsa. The sample information and corresponding accession number for each sample were listed in supplemental Table.

## Acknowledgments

This work was supported by grants from Guangdong Provincial Novel Coronavirus Scientific and Technological Project (2020111107001), Science and Technology Planning Project of Guangdong (2018B020207006).

---

**Supplemental Table.** Sample information and accession no for all sequencing data.

| Sample name | Sequencing method | Cell strain | Passage | Accession NO |
| --- | --- | --- | --- | --- |
| 20SF5645 | PCR+Nanopore | - | original | SAMC150972 |
| ST-N3-D | PCR+Nanopore | - | original | SAMC150973 |
| SZ-N16-D | PCR+Nanopore | - | original | SAMC150974 |
| 20SF014 | Metagenomic | - | original | SAMC151281 |
| 029 | PCR+Nanopore | Vero-E6 | 2 | SAMC150975 |
| 112 | PCR+Nanopore | Vero-E6 | 2 | SAMC150976 |
| 107 | PCR+Nanopore | Vero-E6 | 2 | SAMC150977 |
| 115 | PCR+Nanopore | Vero-E6 | 2 | SAMC150978 |
| 1676 | PCR+Nanopore | Vero-E6 | 2 | SAMC150979 |
| 252 | PCR+Nanopore | Vero-E6 | 2 | SAMC150980 |
| 262 | PCR+Nanopore | Vero-E6 | 2 | SAMC150981 |
| 265 | PCR+Nanopore | Vero-E6 | 2 | SAMC150982 |
| 263 | PCR+Nanopore | Vero-E6 | 2 | SAMC150983 |
| 272 | PCR+Nanopore | Vero-E6 | 2 | SAMC150984 |
| F2 | PCR+Nanopore | Vero-E6 | 2 | SAMC150985 |
| F4 | PCR+Nanopore | Vero-E6 | 2 | SAMC150986 |
| F5 | PCR+Nanopore | Vero-E6 | 2 | SAMC150987 |
| 028 | PCR+Nanopore | Vero | 2 | SAMC150988 |
| 107 | PCR+Nanopore | Vero | 2 | SAMC150989 |
| 115 | PCR+Nanopore | Vero | 2 | SAMC150990 |
| 025 | PCR+Nanopore | Vero-E6 | 2 | SAMC150991 |
| 028 | PCR+Nanopore | Vero-E6 | 2 | SAMC150992 |
| 108 | PCR+Nanopore | Vero-E6 | 2 | SAMC150993 |
| 112 | PCR+Nanopore | Vero | 2 | SAMC150994 |
| 108 | PCR+Nanopore | Vero | 2 | SAMC150995 |
| 014/MiSeq | PCR+MiSeq | Vero-E6 | 3 | SAMC150996 |
| 014/cDNA | Nanopre direct cDNA | Vero-E6 | 3 | SAMC150997 |

